# Extracellular vesicles released from endothelial cells of the blood-brain barrier mediate brain Iron accumulation during LPS-induced brain Inflammation

**DOI:** 10.1101/2025.07.30.667763

**Authors:** Kondaiah Palsa, Timothy B. Helmuth, Aurosman Pappus Sahu, Rebecka O. Serpa, Elizabeth B. Neely, Irina A. Elcheva, Vladimir S Spiegelman, James R. Connor

**Affiliations:** Department of Neurosurgery, Penn State College of Medicine, Hershey, Pennsylvania, USA; Department of Pediatrics, Penn State College of Medicine, Hershey, PA, USA

## Abstract

**Introduction:** Brain inflammation leads to an increase in the amount of iron in brain tissue; however, studies do not address the source of the iron that could lead to the accumulation. Most of the brain iron uptake is mediated through the blood-brain barrier (BBB), but studies have not examined whether inflammation increases or decreases iron flux across the BBB. Our recent in vitro study discovered a novel alternate mechanism that iron transport across the BBB is mediated via the extracellular vesicles (EVs). Herein, we investigated the impact of brain inflammation on iron release and iron transport via EVs from the brain microvasculature (BMV).

**Methods:** For this study, we developed an in vivo brain inflammation model. We induced brain inflammation in three-month-old C57BL/6 by intracerebroventricular injection of lipopolysaccharide (LPS,12μg/mice). For in vitro, we used human blood-brain barrier endothelial cells derived from human-induced pluripotent stem cells (hiPSCs). We separated the BMV from brain parenchyma by using density gradient centrifugation. To inhibit the EVs synthesis, we injected intraperitoneally for 21 days GW4869 (60μg/mice), an inhibitor of neutral sphingomyelinase 2, a key regulatory enzyme necessary for EV formation. The brain EVs were isolated by ultracentrifugation. We measured the BMV and parenchyma iron concentration by Inductively coupled plasma mass spectrometry (ICP-MS). Furthermore, we performed immunoblotting to measure the protein expression in BMV and EVs.

**Results:** The LPS injection activated microglia and astrocytes as well as increased the brain proinflammatory cytokines compared to the control mice. Furthermore, brain inflammation increased the iron levels in the brain parenchyma but decreased the iron levels in BMV. Brain inflammation was associated with the degradation of ferroportin (FPN1), an iron exporter, in the BMV. CD63, an EVs membrane protein, was increased in the BMV and associated with increased FTH1-iron release via EVs from BMVs to the brain. Moreover, brain inflammation induced iron deficiency in BMV as evidenced by an increase in the transferrin receptor and decreased FTH1, suggestive of increased iron uptake. Pharmacological reduction of EVs by GW4869 reduced iron accumulation in the inflamed brain parenchyma compared to control mice.

**Conclusion:** This is the first study to demonstrate that EV inhibition decreases iron in the brain. Degradation of FPN1 in the BMV during inflammation did not limit iron accumulation but there was an increase in FTH1-iron-enriched EVs indicating these are responsible for brain iron accumulation during inflammation. Thus, in summary, we have discovered a novel mechanism that involves BMV-released EVs enriched with iron that is the mechanism for brain iron accumulation during inflammation.

## Introduction

Iron is crucial for various biological functions, particularly in cognition and brain health. It acts as an electron donor, acceptor, and oxygen carrier, making it essential for oxidation-reduction reactions critical to cellular metabolism[1]. While iron benefits the brain, fluctuations in its levels can lead to neurological problems[2]. Blood-brain barrier (BBB) endothelial cells play a key role in iron transport to the brain and thus regulate its availability to the brain[3,4]. The transferrin receptor1(TfR1) on the apical side of endothelial cells in the blood–brain barrier takes up iron bound to transferrin and releases it to the brain parenchyma as free iron via the iron exporter ferroportin1/hephaestin (FPN1/HEPH) or iron-protein complex[3,5]. The FPN1 released iron is regulated by hepcidin, which is released from astrocytes and microglia[6–8]. The release of EVs is iron-regulated by the iron concentration in the endothelial cells of the BBB[9]. However, during inflammation, this process is altered in a way that is not known and addressed in this study.

Inflammation is the primary mechanism for iron accumulation and the progression of neurological disorders[10]. Glial cells, especially microglia, play the main role in inflammation[11]. Under physiological conditions, microglia mainly eliminate metabolic products and toxic materials. Activated microglia, however, increase the release of proinflammatory cytokines, which leads to neuronal death[12]. To induce brain inflammation, studies activate the microglia by using Lipopolysaccharide (LPS), a component that stimulates the immune response via the Toll-like receptor 4 (TLR-4)[13]. TLR-4 is primarily found on microglia in the central nervous system[14] and activation of the microglia produces proinflammatory cytokines such as TNF-α, IL-6, and IL-1β[15]. These cytokines serve as critical mediators of neuroinflammation. LPS administration is frequently employed to investigate diseases associated with neuroinflammation in mouse models[16]. However, the specifics of iron trafficking within blood-brain barrier (BBB) endothelial cells during brain inflammation remain unclear, particularly whether inflammation affects iron transport across the intact BBB.

Our current study aims to investigate the effect of brain inflammation on iron uptake in blood-brain barrier endothelial cells. To this end, we induced brain inflammation through intracerebroventricular (ICV) injection of LPS and examined the iron homeostasis at the brain microvasculature. This study addresses a gap in knowledge regarding the mechanism by which inflammation alters iron flux across the intact BBB.

## Methods

### Animals and treatments

Male C57BL/6J mice, aged 11–12 weeks, were obtained from the Jackson Laboratory. All animal experiments received approval and were conducted following the guidelines of Penn State University’s Animal Ethics Committee(#PROTO201900998). The mice were housed in an environment with automatically regulated temperature (21–25 °C), relative humidity (45–65%), and a light-dark cycle of 12 h. We divided mice into two groups: (I) i.c.v. saline group, or (ii) i.c.v. LPS (12 μg) group, with each group comprising 6 male mice (n=6). The i.c.v. injections of 12 μg LPS (diluted in 3 μL saline) and saline (as a control) were performed on the same day using a microsyringe, following stereotaxic coordinates −2.6 mm dorsal/ventral, −1.5 mm lateral, and −0.2 mm anterior/posterior from bregma as described previously[16].

### EVs inhibition model

GW4869 (N, N’-Bis[4-(4,5-dihydro-1H-imidazol-2-yl) phenyl]-3,3’-p-phenylene-bis-acrylamide dihydrochloride) is the most widely used pharmacological agent to block EVs generation and reduce EVs release by neutral sphingomyelinase (nSMase-2). The nSMase-2 activity is important for creating the large lipid raft domains involved in EVs shedding, as a result, inhibition of nSMase-2 reduces the release of EVs from the cells. GW4869 was prepared and injected as described previously[17–19]. GW4869 was dissolved in DMSO at 5 mg/mL. Mice were injected intraperitoneally with 200 μl of 0.3 mg/mL GW4869 in 0.9% normal saline (60 μg/mouse; 2-2.5 μg/g body weight) or 200 μl of 3.75% DMSO as a saline control every 24 h for four weeks (28 injections total).

We used the same set of animals for PBS and LPS ICV injection study to measure the BMV, parenchyma iron concentration, and immunoblotting, n=6 per group (half of the volume used for each). For EV isolations, we used *n* = 10 per group, n=6 used for immunoblotting, 2 used for the EVs validation, and 2 used for the ferritin iron staining. For the LPS with EVs inhibition group, we used n=10 for each group. n=6 for the LPS injection and 4 were used for the validation of EVs inhibition.

### BBBECs derived from human induced pluripotent stem cells

Human endothelial cells were differentiated from ATCC-DYS0100 hiPSCs (human induced pluripotent stem cells) as described previously[9,20]. hiPSCs were seeded onto a Matrigel-coated plate in E8 medium (Thermo Fisher Scientific# 05990) containing 10 μM ROCK inhibitor (Y-27632, R&D Systems; #1254) at a density of 18000 cells/cm2. The iPSCs differentiation was initiated by changing the E8 medium to E6 medium (Thermo Fisher Scientific A1516401) after 24 h of seeding. E6 medium was changed every 24 h, and cells were continued to E6 medium for up to 4 days. After 4 days, cells were switched to basal endothelial medium (hESFM) (Thermo Fisher Scientific #11111) supplemented with 10 nM bFGF (Fibroblast growth factor, Peprotech # 100– 18B) and 10 μM RA (retinoic acid, Sigma #R2625) and 1% B27 (Thermo Fisher Scientific#17504– 044). The medium was not changed for 48 h. After 48 h, cells were collected and replated onto transwell filters or cell culture plates coated with collagen IV and fibronectin. Twenty-four hours after replating, bFGF and RA were removed from the medium to induce the barrier phenotype.

### Isolation of brain microvasculature

The capillary depletion method was conducted as previously described[21]. Briefly, fresh brain pieces were homogenized on ice by 10 strokes with a Dounce homogenizer (smaller diameter pestle) in 3.5 ml HBSS and then centrifuged for 10 min at 1,000 g. The pellet was resuspended in 2 ml of 17% dextran (mol wt 60,000; 31397; Sigma-Aldrich) and centrifuged for 15 min at 4,122 g to separate the parenchymal cells from the vasculature. The top myelin and parenchymal cell layers were removed together and diluted with HBSS, then centrifuged for 15 min at 4,122 g to pellet the parenchymal cells. Both the vascular pellets and parenchymal cell pellets were resuspended in cold 1% NP40 in PBS with protease and phosphatase inhibitors, agitated for 30 s at 27 Hz with a Tissue Lyser II (85300; Qiagen), and then incubated for 20 min on ice. The cell lysate was collected after centrifugation for 10 min at 12,700 g. The total protein concentration of each sample was measured using a BCA assay.

### EVs enrichment from tissue

Brain EVs isolation was performed as described previously[22]. Modified briefly, brains were sliced on ice using a razor blade to generate 1–2 cm long, 2–3 mm wide sections of brain. The sliced brains were transferred to a 50 mL tube containing 75 U/ml of collagenase type III in Hibernate-E (at a ratio of 800 μL per 100 mg of brain). The tissues were incubated in a shaking water bath at 37°C for a total of 20 min. The dissociated tissue was spun at 300 × g for 5 min at 4°C. The supernatant from the 300 × g spin was then transferred to a fresh tube and spun at 2000 × g for 10 min at 4°C. Finally, the supernatant was spun at 10,000 × g for 30 min at 4°C. The supernatant was overlaid on a triple sucrose cushion (0.6 M, 1.3 M, 2.5 M). To make the triple sucrose cushion, 0.5ml of 2.5 M sucrose (F4) was overlaid with 0.6 ml of 1.3 M sucrose (F3) followed by 0.6 ml of 0.6 M sucrose (F2). The supernatant from the 10,000 × g spin was overlaid onto F2, and the exact volume was recorded (x ml). The gradient was spun for 3 h at 180,000 × g at 4°C. A blank gradient was made and spun at the same time as the sample containing the gradient. After the spin, the top of the gradient was removed and discarded (x ml – 0.6 ml = volume removed), leaving exactly 0.6 ml above F2. This was designated fraction 1 (F1). Fractions 2 and 3 were subsequently collected. Each fraction was diluted with ice-cold PBS and spun at 100,000 × g at 4°C (Beckman Coulter, TLA 100.3) for 1 h to pellet the vesicles. Following spin completion, the supernatant was discarded, and pellets were collected in ice-cold PBS. The vesicles in fractions 2–3 was characterized by nanoparticle tracking analysis, immunoblotting, and TEM.

The EVs isolation from the cell culture media was completed as described by Palsa et.al [9]by sequential centrifugation. Briefly, the basal chamber media were collected and centrifuged at (i) 300g for 10 min to remove the cell pellets, and (ii) the supernatant was centrifuged at 2000g for 10 min to remove the dead cell pellet. The resultant supernatant was centrifuged at 4000g for 30 min to remove the cell debris pellet. The resultant supernatant was collected in a 100K filter (Amicron Ultra-15, Centrifugal filters, Merck Millipore Ltd) to concentrate the media. The concentrated media were collected and centrifuged at 100,000g for 70 min (Beckman Coulter, TLA 100.3). The supernatant was discarded, and the pellet was washed with PBS and again centrifuged at 100,000g for 70 min, with the resultant EV pellet used for downstream processing.

### Transmission electron microscopy

TEM was performed as previously described[9]. Briefly, 10 μl of EVs solution was placed on parafilm. Formvar-coated copper grids were then placed on top of the drops and incubated for 20 min. The copper grids were then incubated with a 4% solution of paraformaldehyde in 0.1 M PBS for 20 min, washed thrice with PBS for 1 min each, incubated with 1% glutaraldehyde in 0.1 M PBS for 5 min, washed with distilled water for 2 min, washed thrice with PBS for 2 min each, contrasted with 1% uranyl acetate for 20 s, and then observed by TEM (JEOL-1400).

### EVs subpopulations isolation

The EV subpopulation was performed as described previously[23]. 400μL of magnetic beads (Invitrogen, MSPB-6003) were washed three times using PBS, and then incubated with Biotin anti-mouse CD54 (1:50, 116103, Biolegend) or, as a control, Biotin Mouse IgG2b, κ Isotype Ctrl Antibody (1:50, 401204, Biolegend) on a shaker at room temperature for 1 h. The beads were then placed on the magnet for 1 min and washed with PBS. 600 μg of EVs (in 0.5ml PBS) was used in the experiment, of which 300μg was incubated with biotinylated anti-IgG antibody or ICAM-1 antibody-coupled magnetic beads at 4°C overnight. After magnet absorption, the supernatant was transferred to a new tube for ultracentrifugation to obtain the EVs after depletion of ICAM^+^ EVs as the total EVs group. The magnetic beads bound with ICAM^+^ EVs were collected and washed 3 times with PBS to obtain purified ICAM^+^ EVs. The same amounts of proteins (12ug) for each group were loaded for western blot analysis.

### Immunoblotting

Immunoblotting was performed as described previously[9]. Briefly, isolated brain microvasculature was homogenized in NP-40 buffer that contained a protease inhibitor cocktail (Cell Signaling Technology, USA) and a phosphatase inhibitor cocktail (Sigma Aldrich, USA). After incubating on ice for 20 minutes, the homogenate was sonicated and centrifuged at 12,000 rpm for 10 minutes at 4°C. The protein content of the supernatant was measured using the micro-BCA kit method. An equal amount of protein (30 μg) was fractionated on 4%–20% SDS-gels under reducing conditions and transblotted onto PVDF membranes. The blots were blocked with 5% fat-free dry milk and probed for CD63 (Thermo Fisher Scientific; 1:1000, 10628D), FPN1(Alpha diagnostic International, 1:1000, MTP-11S), ICAM-1(Invitrogen, 1:1000, MA5407) FTH1 (Cell Signaling Technology, 1:1000, 4393S), TfR1(Santa Cruz Biotechnology; 1:250, sc-65882), tsg 101 (Santa Cruz Biotechnology; 1:250, sc-7964), Calnexin (cell Signaling Technology; 1:1000, 2679), Vinculin (Sigma Millipore, 1:2000, V9131), and beta-actin (Sigma, 1:1000, A5441). A corresponding secondary antibody conjugated to HRP was employed (1:5000, GE Amersham), and bands were visualized using ECL reagents (PerkinElmer) on an Amersham Imager 600 (GE Amersham). The images were quantified with ImageJ software (NIH, Bethesda, MD, USA) and normalized to the respective loading control, β-actin.

### Iron (^57^Fe) Release studies

We have previously described the methodology for performing release assays[20]. Briefly, huECs in serum-free media plated in a 12-well Transwell format were loaded with 100 μM ^57^Fe-ascorbic acid complex and incubated overnight at 37°C. Following incubation, cells and chambers were rinsed three times with 1 × DPBS, and serum-free media was added, along with TNF-α (10 ng/mL) or IL-6 (50ng/mL). These concentrations were selected based on previous studies. ^57^Fe release in EVs was measured after 24h of incubation. As described before, tight junction formation and barrier permeability were measured via TEER and RITC signal.

### Brain iron analysis

Tissue Acid digestion was performed as described previously[24]. Briefly, wet brains were digested overnight with nitric acid (2 mL) and, 30% hydrogen peroxide (1 mL) at 60°C. Iron concentration was determined by ICP–MS against a standard (MERK-multi-element standard # 111355). Results were expressed as μg/g of wet tissue weight.

### Nanoparticle tracking analysis

EVs quantification was performed using a NanoSight NS300 (Malvern Instruments Ltd, Malvern) previously[9]. Briefly, EVs samples were diluted in 1 ml of particle-free water, and each sample was loaded by syringe pump into the NanoSight NS300 (Malvern Instruments Ltd, Malvern) set in scatter mode, and five 60-s videos were generated at 24.98 frames/sec. The size distribution and concentration of particles were calculated, and images were acquired using NanoSight software, version 3.2 (Malvern Instruments Ltd).

### Prussian Blue staining

Prussian Blue staining was performed on the non-denaturing gels to demonstrate that the H-ferritin from the brain EVs contained iron following a procedure described previously[25]. Briefly, whole brain EVs were isolated and heated to 70 degrees centigrade for 5 minutes and loaded onto the non-denaturing gels. Subsequently, non-denaturing gels were stained with a mixture (1:1v/v) of 2% potassium ferrocyanide (K4Fe (CN)6) and 2% 11.6N HCl prepared immediately before use and incubated overnight at room temperature, then rinsed with distilled water until reaching a close-to-neutral pH and clear bands appeared.

### Statistical analysis

All the data were presented as the mean ± SD. Statistical analysis was employed with the GraphPad Prism version 9.0 software. Comparison of differences between two or multiple groups by Student’s t-test and one-way ANOVA respectively. p < .05 was considered statistically significant.

## Results

### ICV injection of LPS activates the microglia and astrocytes

To induce brain inflammation, we injected 12ug of LPS into the brain via ICV[16]. After 48 hours, we perfused the mice with 0.1M PBS, and the brains were collected. The brain was homogenized and separated into BMV and parenchymal fractions. First, we determined the permeability of the BBB injecting the Evans blue dye and measuring the albumin levels in the brain. Our results demonstrated that Evans blue dye was not detected in the brains following ICV injection of either LPS or PBS (**Fig.S1**). Next, we measured the albumin levels in the brain parenchyma and found no significant changes (**Fig.1A**).The GFAP and Iba1 are the markers for the astrocytes and microglia, respectively, whose expression is increased when these cells are activated[16,26]. Therefore, we measured Iba1 and GFAP expression and found the LPS-injected mice have increased expression of GFAP and Iba1 compared to the PBS group (**Fig.1B**). Additionally, we measured the proinflammatory cytokine levels in the parenchyma. The TNF alpha and IL-6 levels were significantly increased in LPS-injected mice compared to the PBS mice (**Fig.1C and D**).

**Fig. 1.**
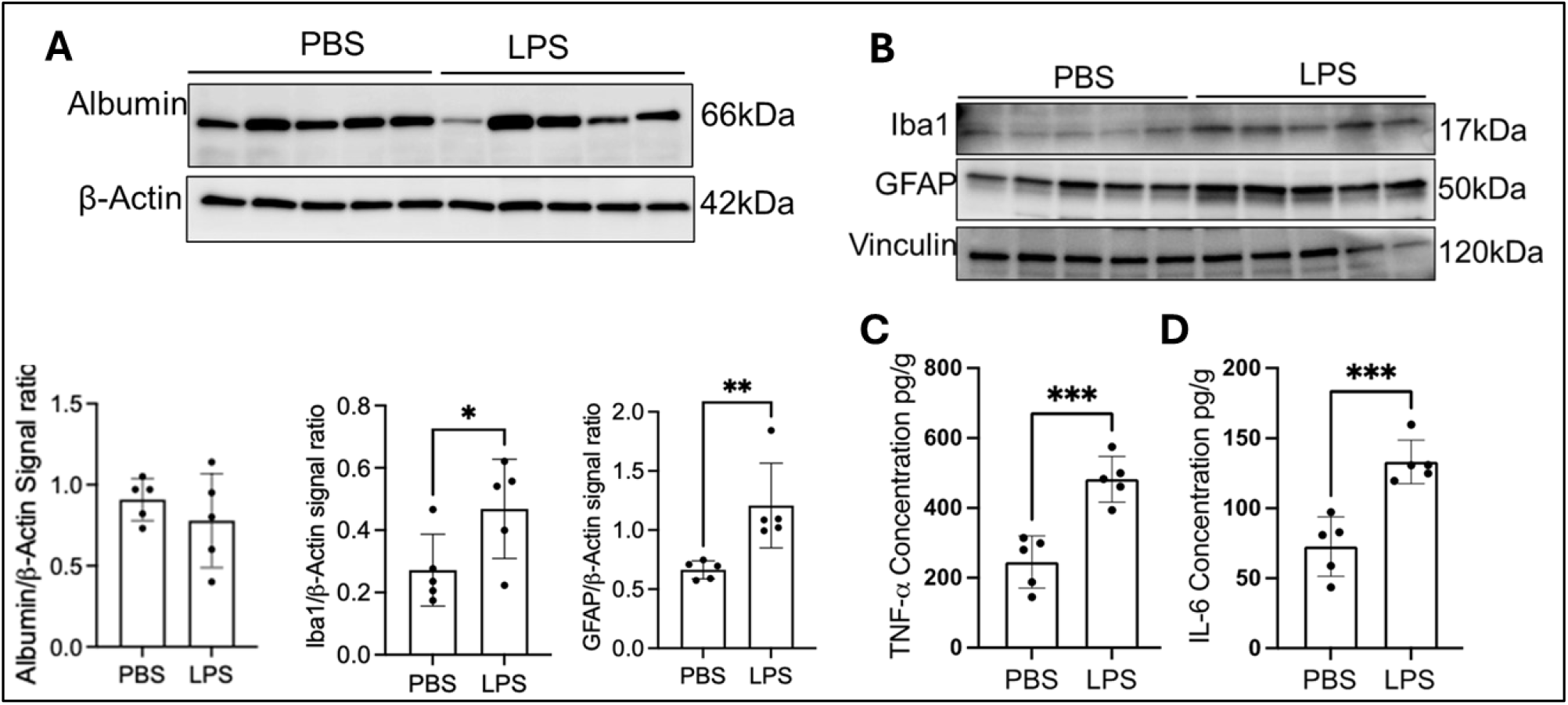
ICV injection of LPS activates the microglia and astrocytes: Twelve-week-old male mice were ICV injected with PBS or LPS and incubated for 48 hours. (A) The immunoblot illustrates parenchyma Albumin levels in PBS and LPS injected groups, and β-Actin is present as the loading control. (B) illustrates immunoblot of the GFAP and Iba1, and β-Actin is present as the loading control. (C) Bar graphs demonstrate the concentration of TNF- and IL-6 levels in the parenchyma of PBS and LPS injected groups. n = 6 for all conditions, the data were presented as mean ± SD, and a student unpaired t-test was used for the statistical significance. *p < 0.05, **p < 0.01, ***p < 0.001

### LPS-induced brain inflammation alters BMV iron, Transferrin Receptor, and Ferroportin

To determine whether brain inflammation affects iron levels in the BMV and parenchyma, we assessed the iron concentration in these areas using ICP-MS. LPS-induced brain inflammation significantly reduced iron levels in the BMV but increased levels in the parenchyma (**Fig. 2A and B**). The iron concentrations in the BMV and parenchyma are dependent on the TfR1 expression. We observed a notable increase in TfR1 expression in the BMV in the LPS group compared to the PBS group (**Fig. 2C**). Additionally, we measured the free iron exporter FPN1 expression, which is significantly lower in the BMV of the LPS group than in the PBS-injected mice (**Fig. 2C**). In summary, LPS-induced brain inflammation results in iron deficiency in the BMV but not in the brain parenchyma.

**Fig. 2.**
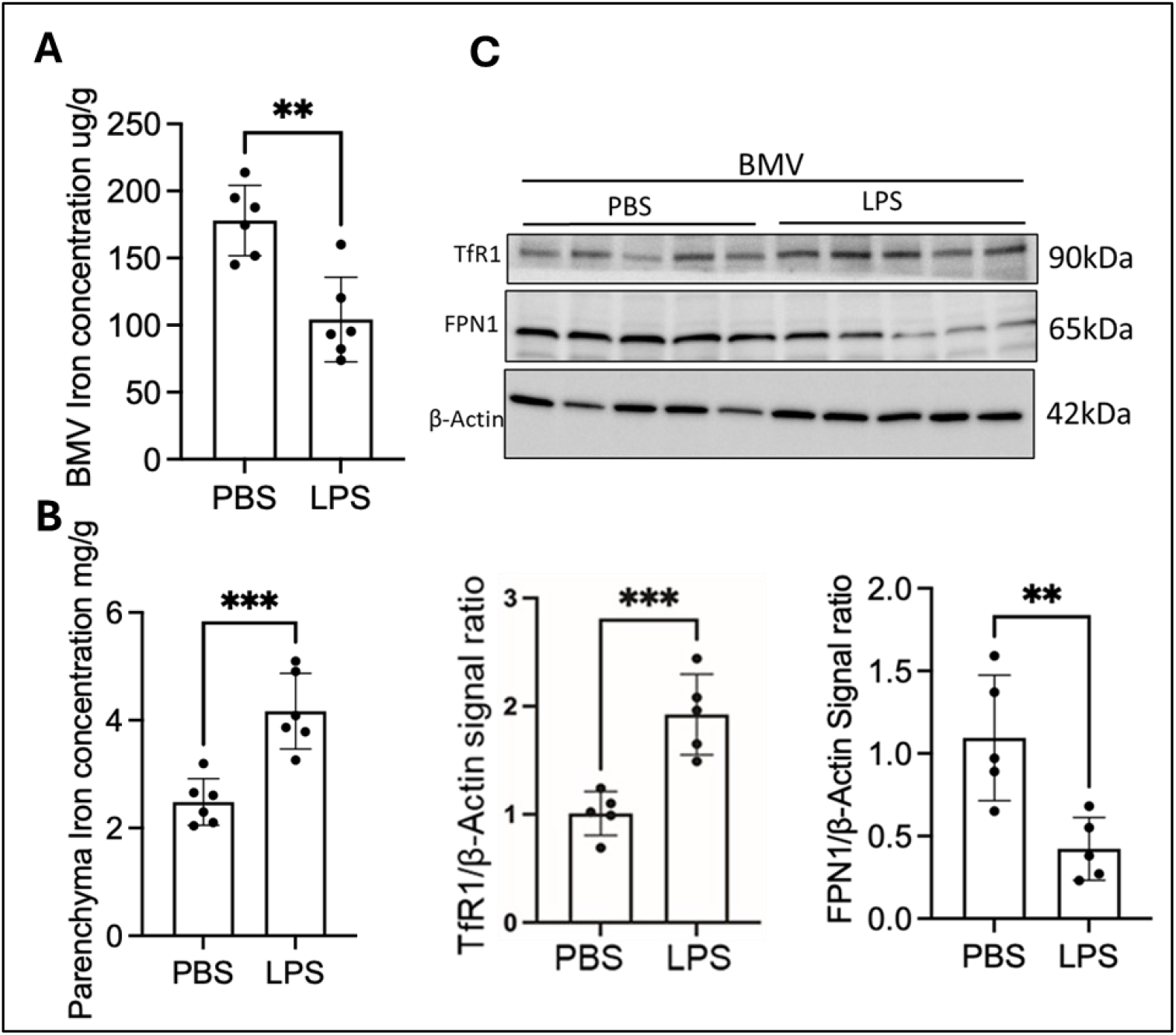
LPS-induced brain inflammation alters the BMV iron and iron uptake protein expressions: Twelve-week-old mice were ICV injected with PBS or LPS and incubated for 48 hours. after that we collected the brain and separated the brain parenchyma and BMV. (A) The bar graph represents the Iron concentration in the BMV. B) Illustrates the iron concentration in the parenchyma. C) Immunoblot and their bar graphs represent the BMV TfR1 and FPN1 expression, β-Actin is present as the loading control. n = 6 for all conditions, the data were presented as mean ± SD, and a student unpaired t-test was used for the statistical significance. **p < 0.01, ***p < 0.001

### EVs membrane protein CD63 expression is increased in BMV during LPS-induced brain inflammation

Our prior research demonstrated that EVs released by ECs mediate FTH1 and iron transport to the brain, and the EV formation is regulated by the iron concentration in ECs[9]. We observed a significant increase in CD63 expression compared to the control group following the LPS injection in the EVs **(Fig. 3A**). Additionally, we measured the FTH1 levels in the BMV of PBS and LPS-injected groups. Brain inflammation was associated with significantly decreased expression of FTH1 in BMV compared to the PBS group (**Fig.3A**). The FTH1 and CD63 expression are regulated by the IRPs in the ECs[9,27]. Thus, we assessed Iron regulatory protein (IRP) expression; brain inflammation reduced IRP2 expression in BMV, but it did not impact IRP1 expression (**Fig. 3C**). These findings indicate that LPS-induced brain inflammation disrupts iron regulatory mechanisms in the BMV; despite lower iron levels, IRP2 expression remained low.

**Fig. 3.**
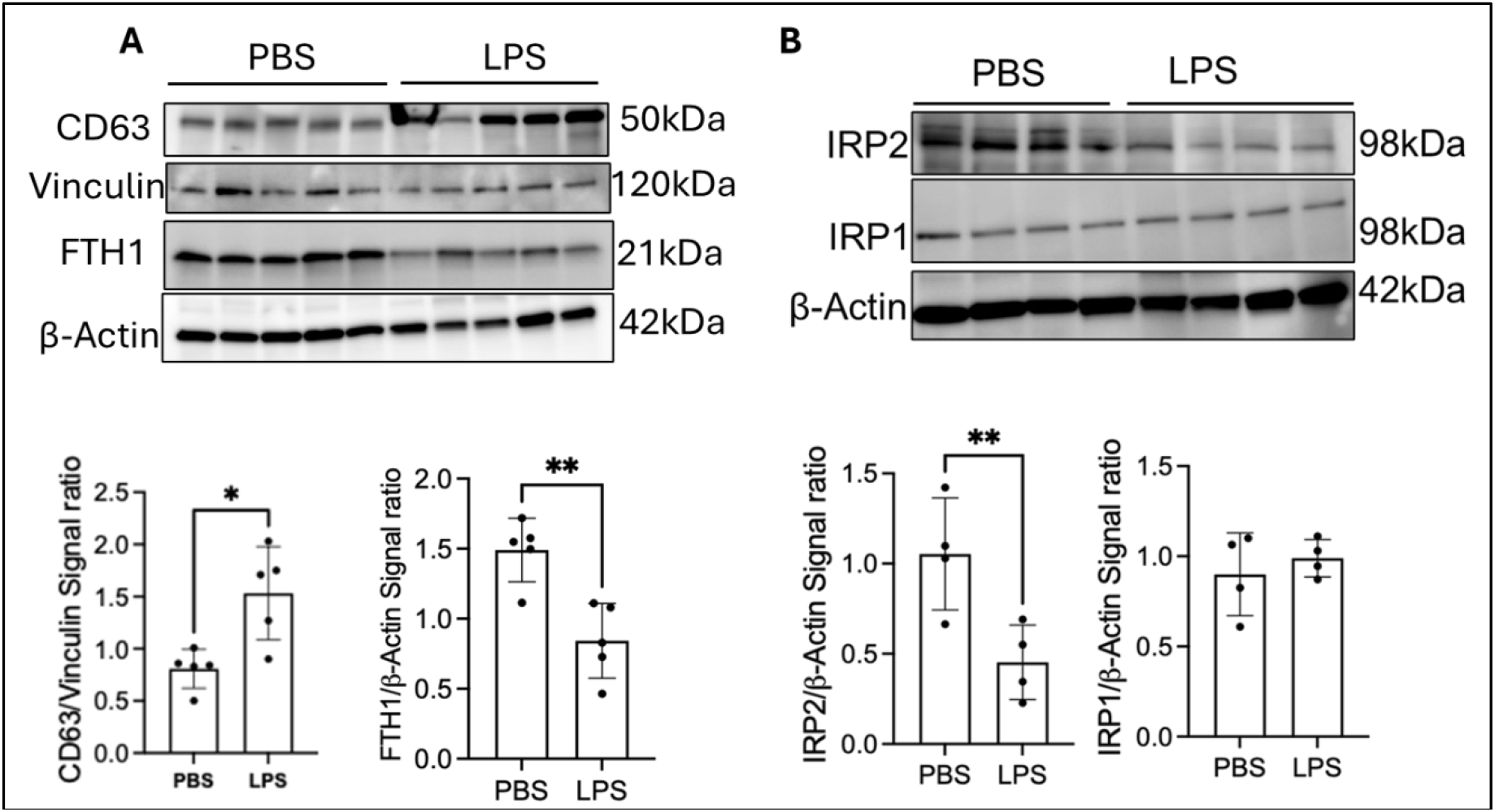
EVs membrane protein CD63 expression increased in BMV during LPS-induced brain inflammation. Twelve-week-old mice were ICV injected with PBS or LPS and incubated for 48 hours. after that, the collected brain is separated into the brain parenchyma and BMV. A). Immunoblot and bar graphs represent the CD63 and FTH1 expression in BMV, and vinculin and β-Actin is present as the loading control, respectively. B) Immunoblot and bar graphs illustrate the IRP 1 and 2 expressions in BMV; β-Actin is present as the loading control. n = 6 for all conditions, the data were presented as mean ± SD, and a student unpaired t-test was used for the statistical significance. *p < 0.05 **p < 0.01,

### LPS-induced brain inflammation increases the release of CD63+EVs from the ECs of the BBB

To determine if brain inflammation is associated with increased release of CD63+EVs from endothelial cells (ECs) and FTH1, we isolated EVs from the whole brain and then assessed the expression of CD63 and FTH1 in the EVs released by ECs. Initially, the EVs were isolated from the normal brain by using density gradient centrifugation. Subsequently, we verified that the isolated particles were indeed EVs by evaluating their size and shape through transmission electron microscopy (TEM), nanoparticle tracking analysis (NTA), as well as checking specific markers via immunoblotting across three fractions. We found that fractions F2 and F3 contained EVs, with F2 having larger EVs, lower concentration, and F3 containing smaller and higher EVs concentration (**Fig.4A, B**). We then analyzed the expression of EV markers and associated proteins—CD63, Tsg101, ICAM-1, and FTH1 within the F2 and F3 fractions (**Fig.4C**). Notably, the levels of EV markers and FTH1 were higher in the F3 fraction, prompting us to utilize this fraction for further studies. Next, we performed Perls Reaction to see whether the EVs associated with FTH1 contain iron. The brain inflammation increased the release of iron-rich FTH1 in EVs compared to the control group (**Fig.4D**). To identify whether the ECs of BBB increase the release of FTH1 in EVs, we focused on ICAM-1, a marker exclusively expressed in on EVs of endothelial cells and increases during inflammation[28]. We isolated ICAM-1+ EVs from the total brain EV population (**Fig.4E**). Here, the control is an IgG Fc antibody added to EVs. Next, we measured the expression of FTH1 and CD63 in ICAM-1+ EVs. Results showed that the expressions of CD63 and FTH1 were significantly higher in the group of mice injected with LPS compared to those injected with PBS (**Fig.4F**).

**Fig. 4.**
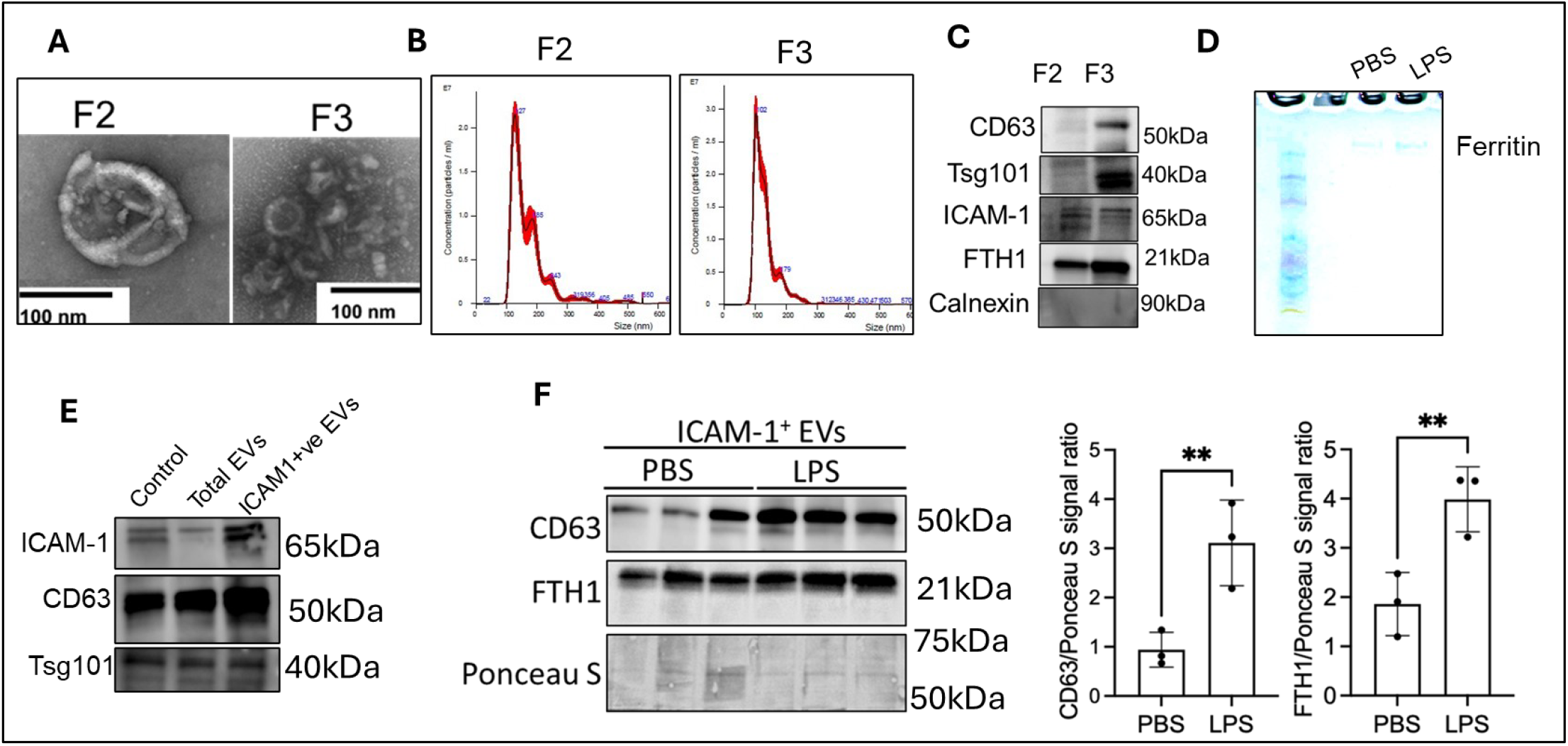
LPS-induced brain inflammation increases the release of CD63+EVs from the ECs of the BBB. Twelve-week-old mice were ICV injected with PBS or LPS and incubated for 48 hours. After that, we isolated EVs from the whole brain by density gradient centrifugation. A) TEM image illustrates the size of the EVs in F2 and F3 fractions. B) Nano particle tracking analysis of F2 and F3 fractions. C), Immunoblot illustrates the EVs markers expression in F2 and F3 fractions-CD63, Tsg101, FTH1, ICAM-1, and calnexin. D) Prussian blue staining of PBS and LPS EVs iron E) Immunoblot represents the ICAM-1+ EVs isolation from the whole EVs. Here, control means the EVs were incubated with IgG2b, κ and whole EVs means ICAM-1 depleted EVs. F). Immunoblot represents the FTH1 and CD63 expression in the ICAM-1+ve EVs and Ponceau S, as used as a loading control. n = 6 for all conditions, each band represents the pooled sample of the n=2, data were presented as mean ± SD, and a student unpaired t-test was used for the statistical significance. **p < 0.01,

### The proinflammatory cytokines increase the iron release from the BBBECs via EVs in vitro

To further confirm ECs of BMV increase the iron release by EVs during brain inflammation, we treated the ECs with proinflammatory cytokines. For this experiment ECs were loaded with ^57^Fe from the apical side and treated with TNF-α (10ng/ml) and IL-6 (50ng/ml) in the basal chamber (**Fig.5A**). 57Fe was added to the apical chamber in the absence of proinflammatory cytokines in the basal chamber as a control. After 24 h, we measured the permeability of ECs by TEER values; the proinflammatory cytokines did not affect the permeability of the ECs at the 24h time point (**Fig.5B**). After measuring TEER values, EVs were isolated from the basal chamber media, and 57Fe and FTH1 activity were measured in EVs. The proinflammatory cytokines significantly increased the 57Fe concentration and FTH1 compared to the control in EVs (**Fig. 5C and D**). Additionally, we measured the FPN and CD63 expression in ECs in inflammatory and control conditions. The exposure to proinflammatory cytokines significantly decreased the FPN1 expression and increased the CD63 levels compared to the control (**Fig.5E**).

**Fig. 5.**
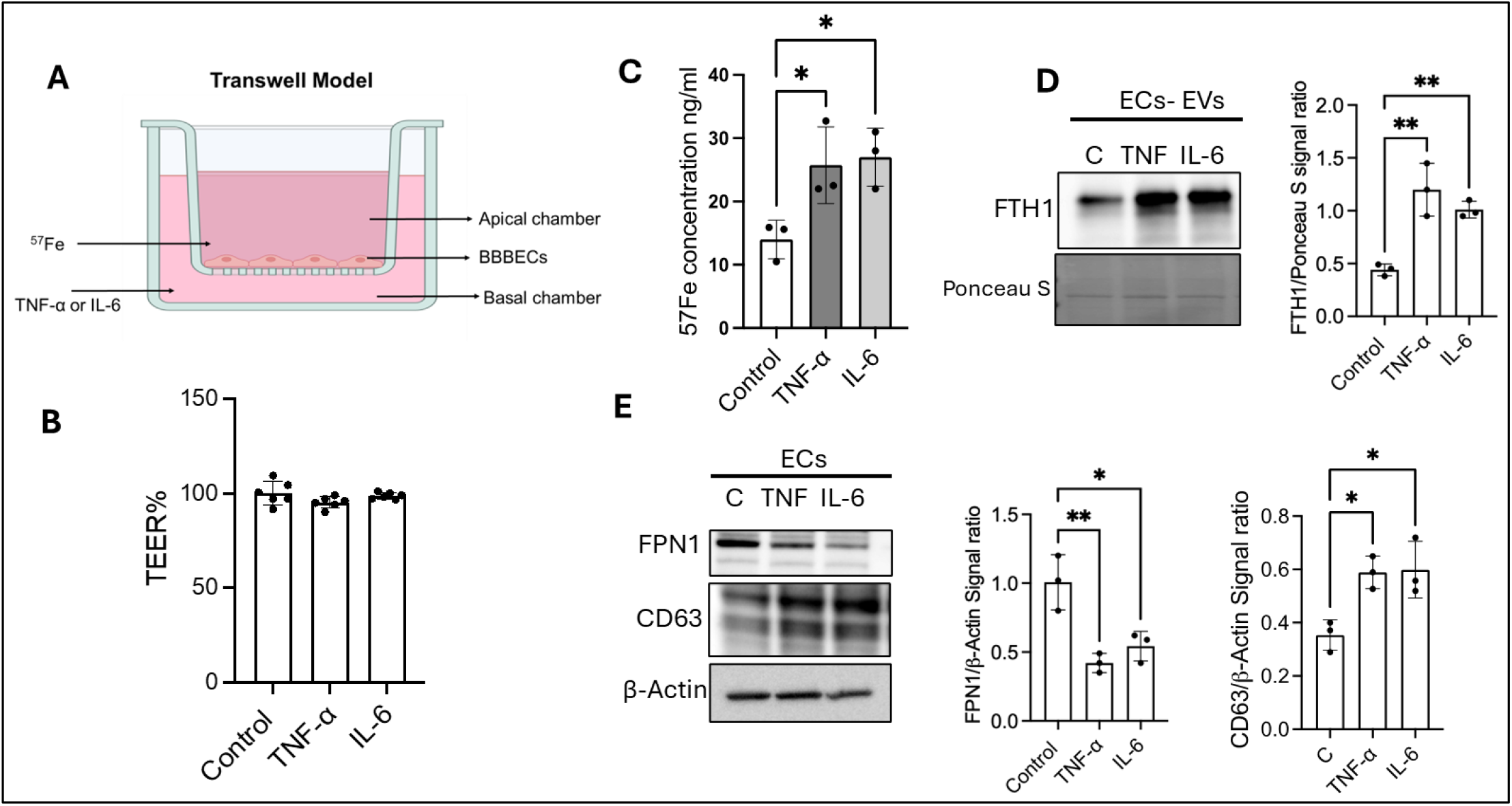
The proinflammatory cytokines increase the iron release from the BBBECs via EVs. ECs were grown on 0.4 μm pore size filters. ^55^Fe-ascorbic acid (100uM) was added to the apical chamber and incubated for 24 hours. After 24h ^57^Fe was removed from the apical chamber, and TNF or IL-6 was added to the basal chamber and incubated for 24h. EVs were isolated from the basal chamber media, and ^55^Fe activity was measured in EVs A) Illustrates the transwell experimental model B) Illustrates the permeability of BBBECs by measuring TEER values in control and proinflammatory cytokines-treated cells. C), bar graph illustrates that 57Fe concentration in control vs proinflammatory cytokines-treated cells. D). The immunoblot and bar graph demonstrate that EVs FTH1 levels. E). Immunoblot and bar graphs demonstrate the FPN1 and CD63 expression in ECs. The experiment was performed in three independent replicates; the data are shown as the mean ± S.D., one-way ANOVA. \**p* < 0.05. **p < 0.01, All experiments

### EVs inhibition by GW4869 reduces the LPS-induced brain iron accumulation

We next inhibited the synthesis of EVs to determine if LPS-induced parenchyma iron accumulation was mediated by EVs. To achieve this, we administered GW4869 intraperitoneally for three weeks (21 days) and then injected LPS via ICV (**Fig. 6.A**). After 48 hours post-LPS injection, we collected and separated the brains into BMV and parenchyma, measuring the iron concentrations. Before LPS injection, we tested whether GW4869 inhibited EVs release into the brain. After 21 days of IP injections, we measured the CD63 expression in whole brain EVs. 21 days of IP injection of GW4869 significantly inhibited the release of EVs to the brain. (**Fig. 6B**). Next, we measured the iron concentration in BMV and parenchyma of EVs inhibited in an inflamed brain. EVs inhibition resulted in increased iron in the BMV group during LPS-induced inflammation compared to those in the LPS alone group (**Fig.6C**). In contrast, the iron level in the parenchyma was significantly decreased when EVs were inhibited during inflammation of the EVs inhibition compared to the LPS group alone (**Fig.6D**).

**Fig. 6.**
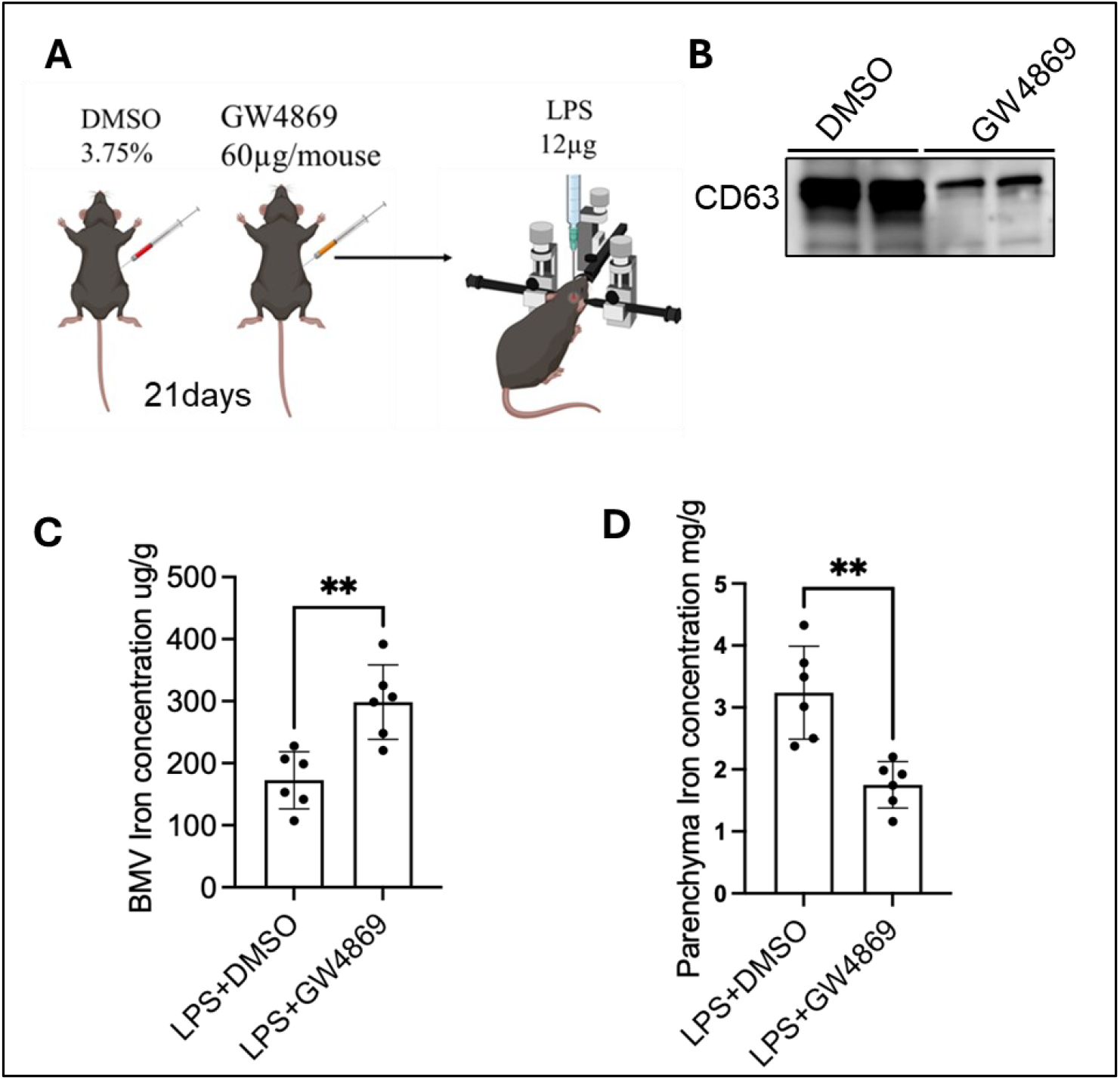
EVs inhibition by GW4869 reduces the LPS-induced brain iron accumulation. Eight-week-old mice were intraperitoneally injected with either DMSO or GW4869 for 3 weeks to inhibit the EVs synthesis in ECs of BMV. After 3 weeks, mice were ICV injected with LPS and incubated for 48 hours; DMSO or GW4869 were continued during and after LPS injections. After that brain was separated into BMV and parenchyma, and the iron concentration was measured by ICP-MS. A). Illustrates experimental design. B). Immunoblot demonstrates the CD63 expression in whole brain EVs, n=2 mice. C) The BMV iron concentration. D) Bar graph demonstrates the parenchyma iron concentration. n = 6 for all conditions, the data were presented as mean ± SD, and a student unpaired t-test was used for the statistical significance. **p < 0.01.

## Discussion

Brain Iron accumulation is a common symptom of neurodegenerative disorders, such as Parkinson’s disease (PD), amyotrophic lateral sclerosis, and Alzheimer’s disease (AD)[29–32]. The potential mechanism regulating this brain iron accumulation is inflammation[33]. In the brain during inflammation, proinflammatory cytokines are elevated, leading to a decrease in FPN1, an iron exporter in EC, and a potential reduction in iron release[34–38]. However, despite this down-regulation of the major iron exporter in the BMV, iron accumulates in the brain. Previous studies showed that iron is also transported from ECs as free and protein-bound iron[4,9]. In this study, we investigated the effects of brain inflammation on BMV iron levels and the expression of their iron transporters. We demonstrated that FTH1 is associated with EVs released from ECs and contains iron. When released during LPS-induced brain inflammation, it could lead to brain iron accumulation. The EV iron release is independent of the hepcidin/FPN1 system. For this study we first confirmed that ICV injection of LPS induces inflammation in the brain. Previous studies showed that microglia in the brain express the LPS receptor; the injected LPS binds to this receptor and activates the microglia[39,40]. The activated microglia release proinflammatory cytokines, which further activate the astrocytes[41]. The current results, consistent with previous studies, demonstrate that ICV injection of LPS activated microglia and astrocytes, increasing the release of proinflammatory cytokines (IL-6 and TNF-α). The activated astrocytes and released proinflammatory cytokines affect the iron homeostasis and regulation in the blood-brain barrier ECs. In the present study, ICV LPS-induced inflammation did not alter BBB permeability for larger molecules, as evidenced by similar albumin levels in both the LPS and PBS injected mouse groups. These results are consistent with a previous study, ICV injection of LPS did not increase the permeability for larger molecules[42].

As previously mentioned, despite the crucial role of endothelial cells in iron transport to the brain, whether inflammation increases or decreases iron flux across the BBB remains unknown. In our current study, we found that FPN1 expression levels in the BMV of LPS-injected mice and BBBECs of proinflammatory cytokines-treated were decreased compared to those in the PBS group, which is consistent with previous studies that LPS induces brain inflammation by releasing proinflammatory cytokines, which degrade FPN1 and elevate TfR1[34,43]. However, our study shows that iron levels were significantly decreased in BMV, which, based on previous studies and ours, would be expected to accumulate due to the iron export protein is degraded. Additionally, the increase in TfR1 expression and decreased FTH1 in the BMV compared to controls suggests that iron uptake is increased and exported through FTH1 as an alternative pathway to FPN to the brain during inflammation. We recently reported that endothelial cells release iron to the brain via extracellular vesicles (EVs) by regulating CD63[9]. In this study, CD63 expression levels in the BMV were found to be higher in the inflamed brain compared to controls; CD63 serves as a marker for EVs, critical for their synthesis and release. CD63 has a 5’ iron-responsive element (IRE) sequence regulated by iron regulatory proteins (IRPs)[27]. The expression of these IRPs depends on cellular iron concentration and is inversely related to iron levels in the cells[44]. Conversely, our study demonstrated a decrease in IRP2 expression, even though the BMV exhibited lower iron levels, FTH1, which indicates that brain inflammation dysregulates the iron homeostasis in ECs of BMV.

Our previous studies showed that FTH1 is involved in the iron transport to the brain during brain development and ID conditions[24,45–47]. Also, FTH1 crosses the blood-brain barrier and is transported intact to the brain cells[48]. During inflammation, the release of ferritin into the circulation increases[49]. Furthermore, studies demonstrated that ferritin (H and L) release in EVs, which are loaded with iron[27,50]. Our in vitro study showed that ECs release the FTH-iron via EVs to the brain[9]. A recent study demonstrated that inflammation increases the release of the iron-rich FTH1 via EVs from the intestinal epithelial cells to the blood. The released EVs’ ferritin iron increases the inflammatory response in macrophages[51]. In the current study, consistent with a previous study, the LPS-induced brain inflammation increased the release of EVs FTH1-iron levels from ECs of BMV in both in vitro and in vivo studies. These EV-associated FTH1 levels were consistent with parenchyma iron levels during LPS-induced brain inflammation. During inflammation, ECs increase the uptake of iron and release FTH1-iron through the EVs to the brain. Additionally, EVs inhibition by GW4869 decreases the iron accumulation in the parenchyma and increases the iron levels in BMV. These EV inhibition results further confirm that ECs of BMV release EVs, mediating the iron accumulation in the brain during inflammation.

Based on our findings, we conclude that LPS-induced brain inflammation disrupts iron homeostasis within the BMV. This disruption in iron homeostasis in the BMV results in overproduction and release of ferritin-enriched EVs from the ECs, resulting in iron accumulation in the brain parenchyma. This is the first demonstration that ECs of the BMV release EVs that are involved in iron accumulation in the brain during inflammation. The mechanism by which the iron accumulation via EVs is occurring may be via the enrichment of FTH1 in the EVs and offers an explanation for how brain iron accumulation can occur during inflammation, by-passing the hepcidin/FPN1 regulatory mechanism. This novel EV-mediated FTH1 transport pathway could provide opportunities to explore mechanisms for the delivery of therapeutic compounds to the brain during the disease process, which typically involves inflammation. Also, these findings suggest potential targets for future clinical applications in patients with iron-accumulated neurodegenerative disease.

## Acknowledgments

We thank Dr Maggie Wang, Ph.D., Laboratory for Isotopes and Metals in the EESI, Penn State University, State College, for the ICP-MS. Drs Han Chen (Manager, Transmission Electron Microscopy Facility, Penn State) and Dr. Jeffrey Sundstrom’s laboratory for assistance with TEM images and NTA, respectively.JRC would like to acknowledge funding received from the National Institute of Health (NIH–R01NS113912). The Fig.5A and 6A were created with BioRender.com.

## Authors contribution

KP and JRC conceived the study. KP, TBH, ROS and APS performed the experiments. EBN supplied reagents and mice. IAE and VSS provided the iPSCs. KP, TBH, ROS and APS analyzed the data. KP and JRC drafted the manuscript. All authors read, edited, and approved the final version of the manuscript.

## Data availability

No datasets were generated or analyzed during the current study.

