## supplemental figure for "Extracellular vesicles released from endothelial cells of the blood-brain barrier mediate brain Iron accumulation during LPS-induced brain Inflammation"

**LPS-induced inflammation did not increase BBB permeability**

Blood-brain barrier (BBB) permeability was assessed using the Evans blue dye method, as previously described[1]. Evans blue is a small azo dye (961 Da) that binds tightly to serum albumin, forming a high-molecular-weight tracer of approximately 69 kDa[2]. A 2% solution of Evans blue in normal saline (4 mL/kg body weight) was administered intraperitoneally to mice that had received either LPS or PBS ICV injections. The dye was allowed to circulate for 24 hours. Following circulation, mice were transcardially perfused with 50 mL of ice-cold PBS, and brain tissues were collected. The results showed that Evans blue dye was not detected in the brains of either PBS- or LPS-injected mice (Figure S.1).

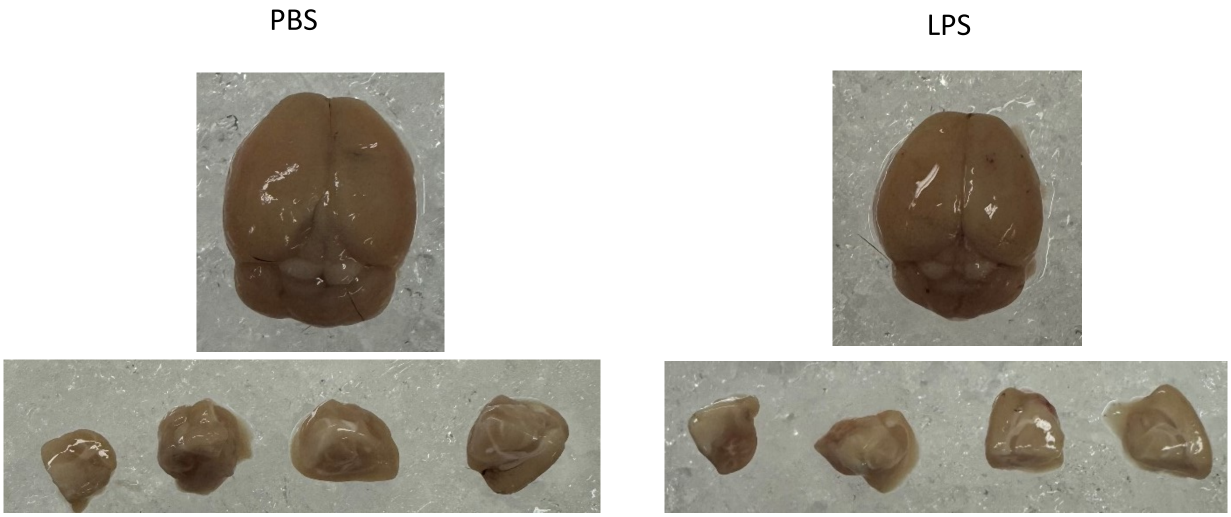


**Fig.S1** LPS-induced inflammation did not increase the permeability of the blood-brain barrier (BBB) to large molecules. Mice injected with either PBS or LPS received an intraperitoneal injection of 2% Evans blue dye and were incubated for 24 hours. After this period, no dye was detected in the brains of either group.
